# SBML to bond graphs: from conversion to composition

**DOI:** 10.1101/2022.05.25.493355

**Authors:** Niloofar Shahidi, Michael Pan, Kenneth Tran, Edmund J Crampin, David P Nickerson

## Abstract

The Systems Biology Markup Language (SBML) is a popular software-independent XML-based format for describing models of biological phenomena. The BioModels Database is the largest online repository of SBML models. Several tools and platforms are available to support the reuse and composition of SBML models. However, these tools do not explicitly assess whether models are physically plausibile or thermodynamically consistent. This often leads to ill-posed models that are physically impossible, impeding the development of realistic complex models in biology. Here, we present a framework that can automatically convert SBML models into bond graphs, which imposes energy conservation laws on these models. The new bond graph models are easily mergeable, resulting in physically plausible coupled models. We illustrate this by automatically converting and coupling a model of pyruvate distribution to a model of the pentose phosphate pathway.

**Graphical Abstract:** 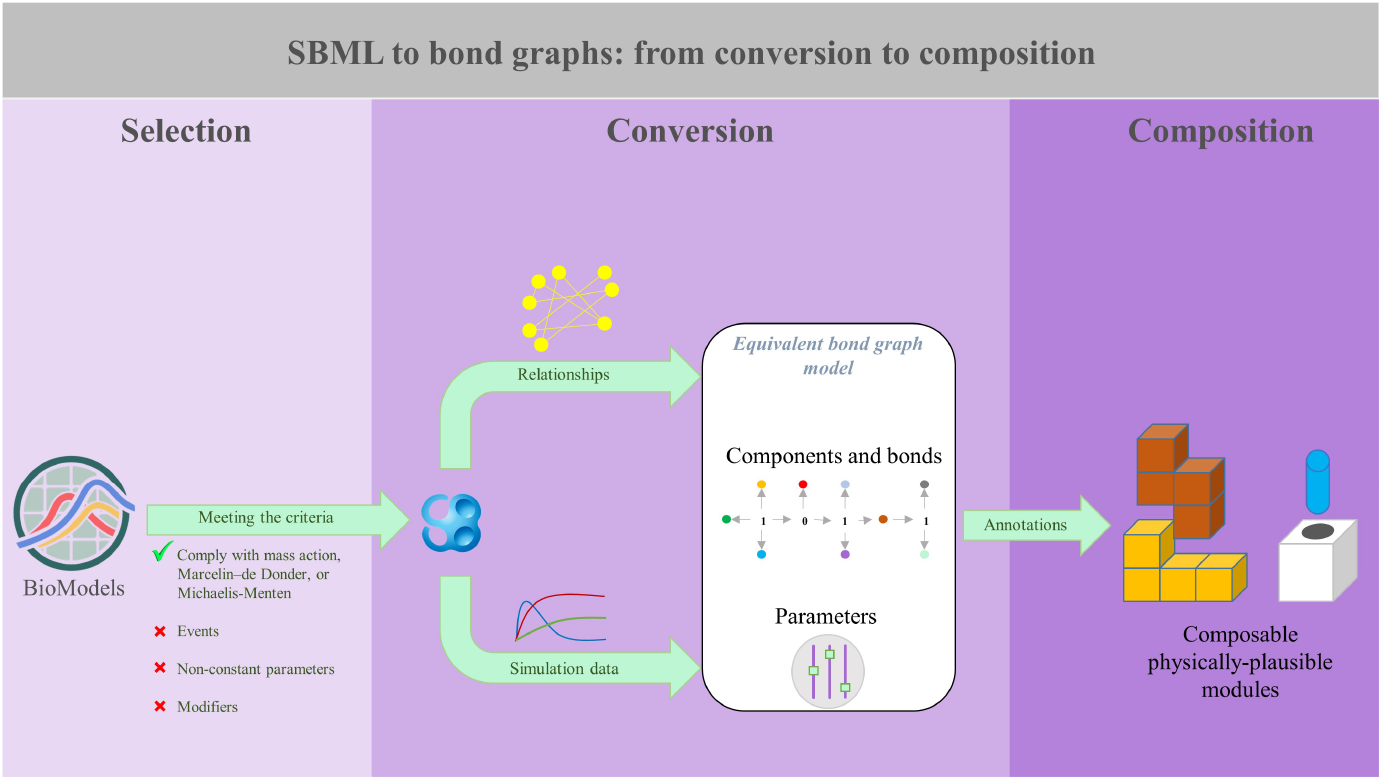

**Highlights:**

- A framework to convert suitable SBML models of biochemical networks into bond graphs is developed.
- The framework is applied here to two interconnecting models of metabolism pathways.
- We automatically integrate the generated bond graph modules.
- We qualitatively illustrate the functionality of the composed model.

## 1. Introduction

Complex biological systems incorporate nested layers of components and interactions which are mostly believed to be naturally organised hierarchically [1, 2, 3]. Accordingly, to model such complex systems, we need to reuse models in a hierarchical fashion [4]. Hierarchical modelling is the composition of existing smaller models, *modules*, where each module can be tested and operated individually. This approach facilitates large-scale model composition by reducing possibilities for human error. In recent years, the Physiome (www.physiomeproject.org) and Virtual Physiological Human (VPH) (www.vph-institute.org) projects have taken initial steps to construct more realistic models to describe systems in the body using modular and hierarchical model development [5, 6].

Biosimulation models are accessible on public repositories such as the Physiome Model Repository (PMR) [7] and BioModels [8], which store models in XML-based formats such as CellML [9] and SBML [10, 11]. However, the available models often violate the laws of physics and thermodynamics, which must be obeyed for any model to be physically realistic. It is clear that combining physically impossible models leads to unrealistic composed models which cannot be used in real-world applications with any level of confidence. Here, we use a framework through which suitable SBML models of biochemical networks can be converted into a modular and physically consistent format to support model composition.

Just like in the natural world, energy is conserved in biophysical processes, regardless of whether they are chemical, mechanical or electrical [12]. Using the principle of energy conservation in the models ensures that all the individual models remain thermodynamically and physically consistent [13, 14]. The Bond Graph (BG) paradigm is an energy and physics-based modelling framework that supports modular and hierarchical modelling.

Bond graphs were invented by Henry Paynter and were primarily meant to be used in mechanical systems [15]. Bond graphs intrinsically follow the energy conservation laws and generate dynamic models based on the laws of physics and thermodynamics. The application of bond graphs was extended to the chemical domain by Borutzky et al. [16] and later by Cellier [17], and to biophysical systems by Oster et al. [18, 19]. More recently, Gawthrop and Crampin developed the application of bond graphs in modelling biochemical and electro-chemical systems [14, 20]. In this paper, we use bond graphs to generate thermodynamically consistent versions of existing biochemical SBML models.

Semantic annotations add a layer of standard biological knowledge as metadata to models to avoid misleading or incorrect naming by modellers. Annotating the models makes them reusable, either solely or in combination with other models [6]. In computational models of biology, modellers are encouraged to label the mathematical content of their models with semantic annotations instead of choosing arbitrary names. This complies with the FAIR data principles (Findable, Accessible, Interoperable, and Reusable) which leads to unambiguous merging of common entities in model composition [21, 22].

Tools and software have been developed for SBML model composition such as the SBML Hierarchical Model Composition (SHMC) package. The SHMC package enables hierarchical modelling of SBML models and supports appending, deleting, replacing, and modifying models’ elements including species, units, and rate laws. However, to ensure compliance with the laws of physics, the model composition process requires changes in equations at points where common species have been detected across the modules [23]. In the SHMC package, such changes in equations are not applied automatically and difficult to write manually. To address this issue, we employed our previously developed framework using bond graphs to compose CellML models and modified it for use with SBML models [24].

In this paper, we focus on composing SBML models that represent biochemical networks. Here, we present an illustrative example in which, using our developed tool, we have automatically converted two SBML models of cellular respiration metabolism into bond graphs: a model of glycolysis and pyruvate metabolism, and a model of the pentose phosphate pathway (PPP) and the tricarboxylic acid (TCA) cycle. Henceforth, we refer to the former as the *pyruvate distribution* model and the latter as the *PPP* model. We have integrated these models into one composite model using our previously developed method in [24].

In this paper, we explain how biochemical reactions are generally expressed in bond graphs (Section 2.1) and how different types of biochemical reactions are converted into bond graphs (Section 2.2). We introduce the overall workflow of our developed model conversion framework to produce bond graph equivalents of SBML models in Section 2.3. To demonstrate the application of our method, we utilised our framework to automatically convert two exemplar SBML models into bond graphs (Section 2.4). Next, we verify and compare the behaviours of the original and bond graph models individually and integratively in Section 3. Finally, remarks of our framework, shortcomings, and the future developments are outlined in Section 4.

## 2. Materials and Methods

In this section, we give a brief introduction to general bond graph modelling of biochemical reactions and how existing formulations of rate laws can be expressed in bond graphs. Based on this, we introduce the workflow of our framework which integrates the bond graph conversion methods of biochemical reactions and automatically implements the required modifications. Our framework is applicable to single SBML models or multiple SBML models in composition. We demonstrate this by converting two SBML models in single mode and in composition.

### 2.1. Biochemical reactions in bond graphs

Here, we explain the fundamental bond graph elements in modelling biochemical reactions and elucidate the equations for single and linked reactions with an example.

Bond graphs are a domain-independent modelling approach that explicitly describes the bidirectional flow of energy between the bond graph elements. Bond graph elements consist of two categories: components and junctions. In biochemistry, the primary components are species, reactions, stoichiometries, sources of flow and potential. Junctions define the conservation laws of the system. At a 0 : *u* junction, all the chemical potentials are equal and all the molar flows sum to zero. At a 1 : *v* junction, all the molar flows are equal and all the chemical potentials sum to zero. Energy is the product of potential (*u*) and flow (*v*) over time: *E* = ∫ *uv dt*, travelling bidirectionally between components and junctions through bonds (shown by ⇀). For further details regarding the application of bond graphs in modelling biochemical systems, we refer the reader to the works by Gawthrop & Crampin [25, 26].

Biochemical processes can be represented in bond graphs using the elements described in Table 1.

**Table 1.** Bond graph elements for modelling biochemical processes.

|  | nature | definition | constitutive relation |
| --- | --- | --- | --- |
| $C_e$ | component | species | $u_x = RT \ln K_x \cdot X$ |
| $C_S$ | component | chemostat (source of chemical potential) | $u_{x_s} = RT \ln K_{x_s} \cdot X_s$ |
| $S_f$ | component | source of flow | $v = f$ |
| $TF : N$ | component | stoichiometry: stoichiometric coefficient | |
| $Re$ | component | reaction | $v = r(e^{u_1/(RT)} - e^{u_2/(RT)})$ |
| $0 : u$ | junction | common chemical potential | |
| $1 : v$ | junction | common molar flow | |

$u$ : chemical potential, $v$ : molar flow, $R$ : ideal gas constant, $T$ : temperature, $X$ : molar concentration, $K_x$ : species constant, $X_s$ : chemostat molar concentration, $K_{x_s}$ : chemostat constant, $f$ : constant inward flow, $r$ : reaction rate constant.

*K_x_* is the species constant and corresponds to the kinetic free energy of a species to participate in reactions and is defined as 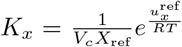 where *V_c_* is the volume of the compartment, *X*_ref_ is the reference concentration (normally 1 mol), and 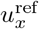 is the standard free energy formation of the species *x* [27]. Chemostats (*C_S_*) are species with fixed concentrations that are considered as sources of potential in bond graphs [20, 28]. Similarly, when there is a constant flow of a species entering the system, we represent it with a source of flow (*S_f_*) in bond graphs. This process is typically called *synthesis* in SBML models. The complete disappearance of a species or physical entity is usually called *degradation* in SBML models. We modelled the process by adding an auxiliary *C_e_* component as the product of degradation with a 1000 times smaller *K_x_* compared to its undegraded reactant. This is because degradation is formulated similar to an irreversible reaction where the product is not specified. We will further discuss the representation of irreversible reactions in bond graphs in Section 2.2.2.

The Boltzmann’s formula is the constitutive relation for the species and chemostats, and the constitutive relation for reactions is the Marcelin–de Donder equation. By substituting the chemical potentials in the Marcelin–de Donder equation with the Boltzmann’s formula for the reaction *X*_1_ ⇌ *X*_2_, we have:

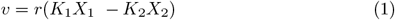

which can also be formulated in mass action kinetics:

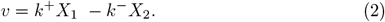

where *k*^+^ and *k*^−^ are the forward and reverse kinetic rate constants in reversible mass action kinetics. Constitutive equations for reactions in bond graphs are defined using the Marcelin–de Donder equation, requiring the species constants and reaction rate constants (bond graph parameters) to be separately defined.

A simple example of two reactions in bond graphs is illustrated in the upper panel in Fig 1. Note that the species B is present in both reactions. The lower panel in Fig 1 demonstrates the composition of Reactions I and II and how the conservation equation changes at the merging point in bond graphs.

**Fig 1.**
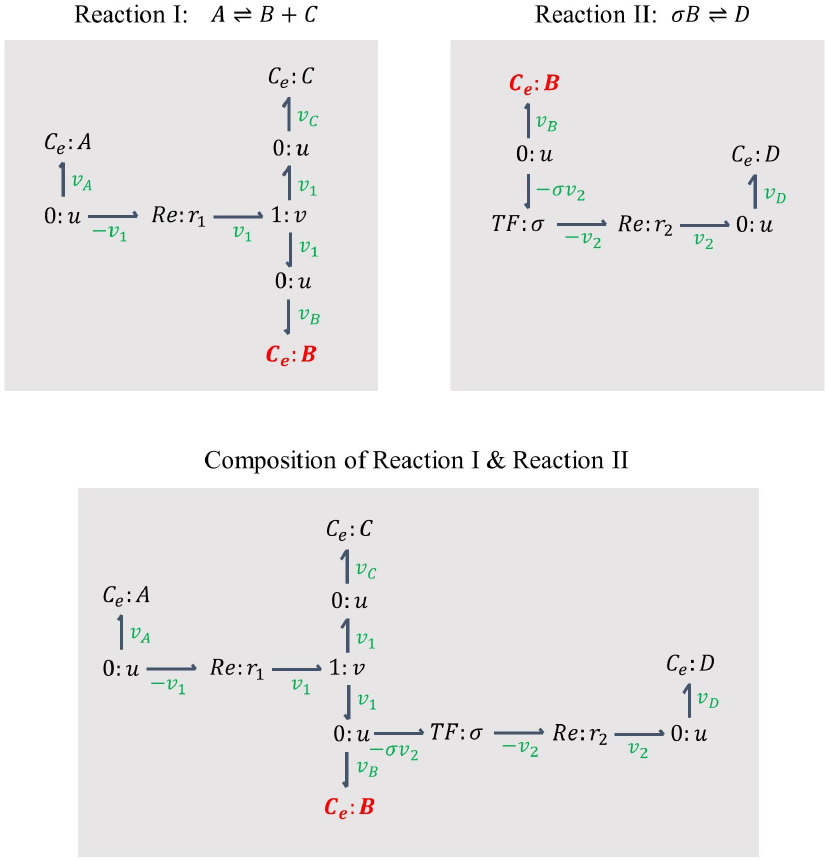
Two exemplar reactions and their composition in bond graphs. Reactions I and II represent two separate reactions in which the species B is common. In composition, the common species (B) is merged and the conservation equation at its corresponding ‘0 : *u*’ junction alters to account for the imposed changes in structure. The conservation equation at the ‘0 : *u*’ junction connected to the species B is *v_B_* = *v*_1_ in Reaction I and in Reaction II is *v_B_* = –*σv*_2_ and in the composed reaction it changes to *v_B_* = *v*_1_ – *σv*_2_.

The reaction rates for Reactions I and II are:

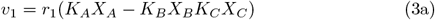

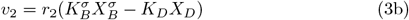

where *r*_1_ and *r*_2_ are the reaction rate constants for Reactions I and II, respectively, *K_A_, K_B_*, *K_C_, K_D_* are the species constants for A, B, C, D, and *X_A_, X_B_*, *X_C_*, *X_D_* are the concentrations of A, B, C, D. *σ* is the stoichiometric coefficient for B in Reaction II. By having the reaction rates, the conservation laws at the ‘0 : *u*’ junctions connected to the species in Reaction I would be as:

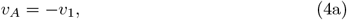

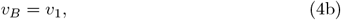

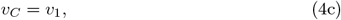

and at the ‘1 : *v*’ junction would be:

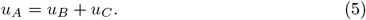

The conservation laws at the ‘0 : *u*’ junctions connected to the species in Reaction II would be as:

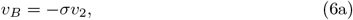

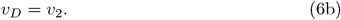

The molar flow rates for the species in the case of composing Reaction I and Reaction II would be defined as:

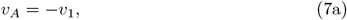

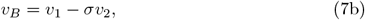

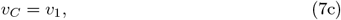

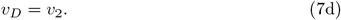

The conservation law at a junction changes if a bond is connected or removed. Here, at the ‘0 : *u*’ junction connected to B in the lower panel of Fig 1, the conservation law demands all the molar flows sum to zero, hence, the attachment of another bond changes the molar flow equation for B (compare Eqs 4b and 6b with Eq 7b).

Utilising the bond graph elements in the following section, we illustrate the procedure of converting existing biochemical reactions into bond graphs.

### 2.2. Bond graph conversion

This section combines the bond graph principles described in Section 2.1 with the extracted data from SBML models to create their bond graph equivalent structures.

Biochemical reactions are constrained by the laws of thermodynamics and can only advance in the direction of decreasing the chemical potential [14]. In general, irreversible reactions are not thermodynamically plausible. Thus, all bond graph models of biochemistry represent reversible reactions. In the next section, we show how reversible and irreversible reactions are described in a reversible fashion in bond graphs.

Briefly, the parameters are estimated by solving equations based on the equilibrium constants (concentration of all products over concentration of all substrates at equilibrium) [29]. The methodology solves for the species parameters prior to finding the reaction parameters. These constraints can be achieved by relating the bond graph parameters to equilibrium constants. This results in a linear matrix equation relating the log-transformed parameters in the general form of:

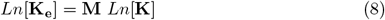

where **K**_*e*_ represents the equilibrium constants, **M** represents the stoichiometric matrix, and **K** represents the unknown species constants. A function in our framework solves this matrix equation and finds the solution with the least square error for the species constants. When bond graph models are composed together, we combine all constraints into a single matrix equation to deal with potential inconsistencies between models. Eq 9 shows the form of the matrix equation in model composition.

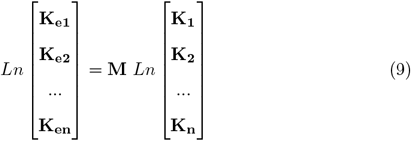

where **K_e1_**, …, **K_en_** represent the equilibrium constants for **n** models and **K_1_**, …, **K_n_** represent the unknown species constants for **n** models.

Once the species constants have been determined, the reaction rate constants are calculated. If the kinetic constants of a model are properly annotated and distinguishable, the reaction rate constants can be calculated directly from the linear matrix; otherwise, they will be achieved by fitting the simulation data to the bond graph equations for reactions. We describe these methods in more detail below.

#### 2.2.1. Reversible reactions

Reversible reactions inherently comply with the bond graph constraints. If a reaction is described using the Marcelin–de Donder kinetics, bond graph parameters can be directly derived. Bond graph parameters need to be estimated in any other case. Here, we discuss how our framework converts the reversible mass action and Michaelis-Menten kinetics into bond graphs.

- **Reversible mass action:** Fig 2.A demonstrates how a reversible mass action reaction is represented in bond graph scheme. Eq 10 describes an exemplar reversible reaction using the mass action kinetics where *k*^+^ and *k*^−^ are the forward and reverse kinetic rate constants, {*X_S_i__* | *i* ∈ {1, 2}} are the concentrations of substrates, {*X_P_j__* | *j* ∈ {1, 2}} are the concentrations of products, and *α* and *β* represent the stoichiometric coefficients.

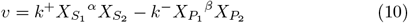 When the bond graph structure of the reactions is determined, we need to find the bond graph parameters. We might face two cases where *k*^+^ and *k*^−^ are properly annotated and distinguishable, or otherwise. In the former case, the bond graph parameters (reaction rate constants and species constants) are best estimated by log-transforming and solving a system of equations for *k*^+^ and *k*^−^ (Eqs 11a and 11b).

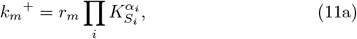

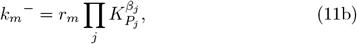

where *m* is the number of reactions [30]. In cases where *k*^+^ and *k*^−^ are not annotated, they are not recognisable and we need to use the simulation data. To estimate the bond graph parameters from the simulation data, the amount for the flux and concentrations at two separate times are extracted (here we have selected initial and final points denoted by 0 and ∞, respectively) and inserted in the equations Eqs 12a, 12b. By dividing the equations at the two points, the ratio between the species constants of the product(s) and substrate(s) will be achieved (Eq 13). *γ* is the value gained from dividing the amounts for the initial and final flux values (Eq 12c). Eq 13 provides a constraint on selecting values for the species constants.

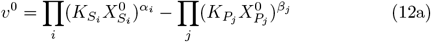

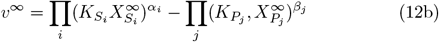

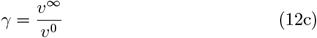

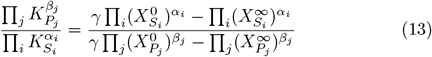 An example of applying both approaches to a system of two interconnected reactions is given in Appendix A. By taking logarithms on each side of the constraint (Eq 13), the relationship between the bond graph parameters can be expressed linearly. By applying this method to all the extracted constraints from a model’s reactions, we generate a linear matrix of constraints. Once solved, the estimated values of the species constants can be used to calculate the reaction rate constants. We achieved this by fitting the simulation data to the Marcelin–de Donder reaction kinetics.
- **Reversible Michaelis-Menten:** In the reversible Michaelis-Menten kinetics a substrate binds with an enzyme to form a complex. The complex then goes through a second reaction to form a product [31]. To convert a reversible Michaelis-Menten reaction into bond graphs, an enzyme catalysed reaction scheme was taken. Fig 2.B illustrates the bond graph equivalent and representation of an exemplar reversible Michaelis-Menten kinetics. The reaction rate law for the reversible Michaelis-Menten can be formulated as Eq 14 ([31, 32]):

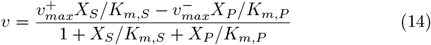

where:

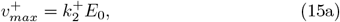

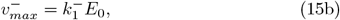

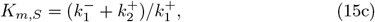

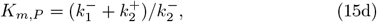

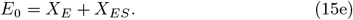

*X*_E_ and *X*_ES_ are the concentrations of E and ES, respectively, and *E*_0_ is the total concentration of the E and ES. By replacing the Marcelin–de Donder parameters in Eq 14 we reach a formula with bond graph parameters [20]:

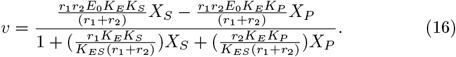 Since normally the equation constants in SBML models are not annotated, we cannot directly extract the original Michaelis-Menten constants. As an alternative, we have access to the simulation data (reaction rates and concentrations) and we can fit the data (*X*_S_, *X*_P_, and *v*) to Eq 16. This will give us the values for four constants where:

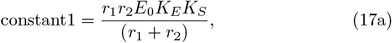

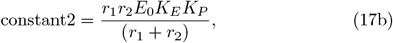

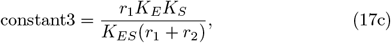

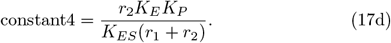 Dividing Eq 17a and Eq 17b gives the ratio between the species constants of reactant and product (*K*_S_ and *K*_P_) and dividing Eq 17c and Eq 17d reveals the correlation between the species constants and the reaction rates (*r*_1_ and *r*_2_). The log-transformation of these two relationships will be added to the linear matrix of constraints. Once the matrix is solved and the amounts for *K*_S_, *K*_P_, *r*_1_ and *r*_2_ are calculated, our framework finds *K*_E_ and *K*_ES_ by setting *E*_0_ = 1 (see Appendix B). An example of converting an SBML model containing two interacting reversible reactions (mass action and Michaelis-Menten) to bond graphs is deposited on GitHub: [Reversible mass action and Michaelis-Menten](https://github.com/Niloofar-Sh/SBMLtoBGs/blob/main/reversibleMA_reversibleMM_irreversibleMA_irreversibleMM/reversMAMM.ipynb).

**Fig 2.**
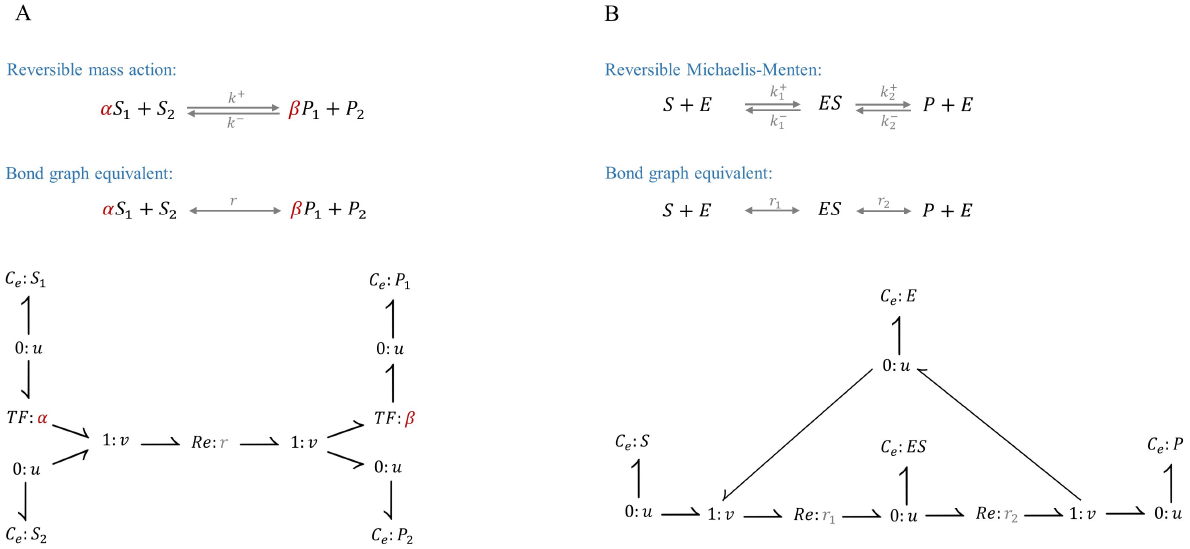
Bond graph representation of reversible reactions. (A) Reversible mass action. *a* and *β* represent the stoichiometric coefficients which are modelled with a transformer in bond graphs; (B) Reversible Michaelis-Menten.

#### 2.2.2. Irreversible reactions

Our framework specifically covers the conversion of two types of irreversible reactions: Irreversible mass action and irreversible Michaelis-Menten. As discussed in Section 2.2.1, any other irreversible format will be treated as an irreversible mass action. Fig 3 demonstrates the bond graph representation of irreversible mass action and Michaelis-Menten kinetics.

**Fig 3.**
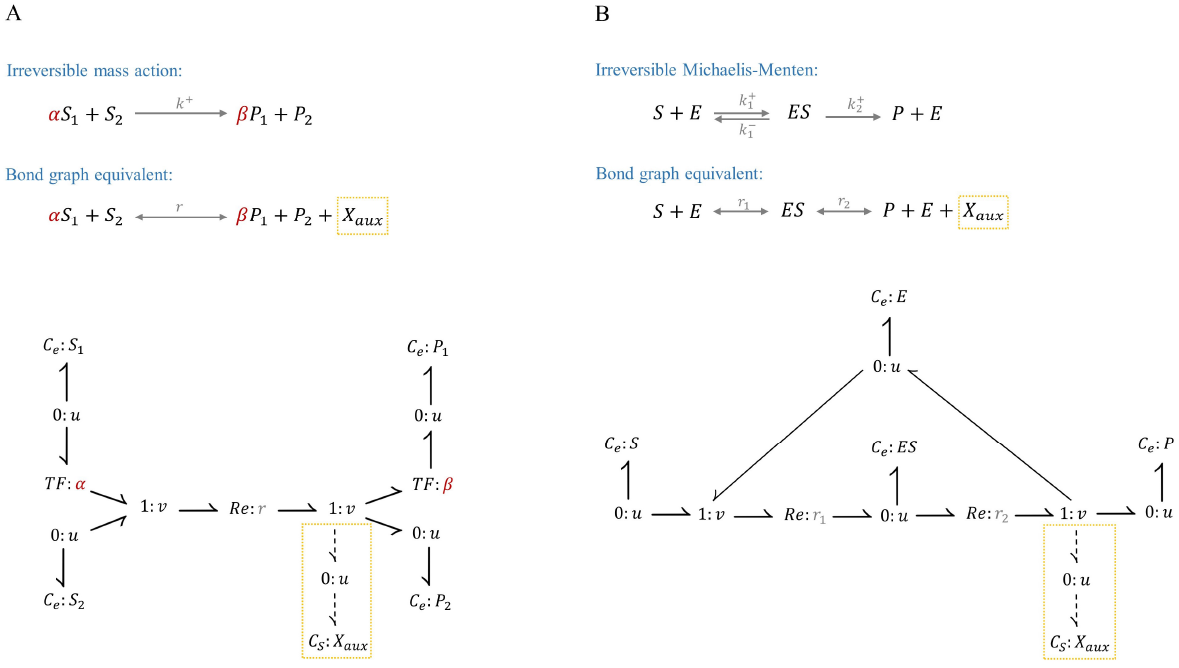
Bond graph representation of two irreversible reactions. (A) Irreversible mass action. *α* and *β* represent the stoichiometric coefficients which are modelled with a transformer in bond graphs; (B) Irreversible Michaelis-Menten. *C_S_* : *X_aux_* is an auxiliary species added to the right side of irreversible reactions in case of thermodynamically inconsistent biochemical loops. Here, the auxiliary species have small *K*s relative to the reactants’.

To approximate irreversibility, the bond graph structures in Fig 3 include auxiliary species (with relatively small *K*s) at the product(s) side of the irreversible reactions. This approach assumes that all irreversible reactions are supplied by an energy from the auxiliary species. Physiologically, adding such an auxiliary species to an irreversible reaction plays the role of missing energy sources which essentially push the reaction forward and hamper it in reverse direction. Hence if the energy providers stop working, the energetically linked reactions will be terminated. Examples of such associated sources of energy would be ATP-ADP-Pi [33, 34, 35], and NADH-NAD [36].

The function of auxiliary species is to avoid miscalculations in the case of thermodynamically inconsistent biochemical loops. In this way, the auxiliary species’ constants handle the additional constraint of representing the irreversible reactions with reversible ones (selecting small values for *K*s at the right side of the reaction). In the following section, we will further explain the selection of *K*s in irreversible reactions.

- **Irreversible mass action:** An irreversible reaction expressed in mass action kinetics is convertible into bond graphs in the same way as its reversible version, except that we assume that the forward flux is far greater than the reverse flux (Eq 18).

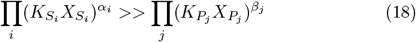 Considering the whole range of concentrations across the simulations, the constraint in Eq 18 can yield:

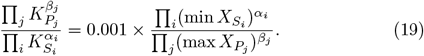

where **min** and **max** represent the minimum and maximum concentrations that a species reaches across the time course of the simulation data. We added the 0.001 coefficient to confine the reverse flux by limiting the *K* values of the products to replicate the irreversibility of reactions. Eq 19 satisfies the law of reversibility of reactions in bond graphs while maintains the pseudo-irreversibility of the original reactions by limiting the reverse flux. While this method applies to any irreversible reaction described in mass action kinetics, it fails to thoroughly capture the behaviour of such reactions if they associate with thermodynamically inconsistent loops. The estimation of bond graph parameters from the kinetic parameters becomes more inaccurate if the species have multiple roles in different reactions, *i.e*., not all the constraints can be satisfied. To best satisfy the constraints in such networks, our framework provides an option of adding an auxiliary species (*C_S_* : *X*_aux_) to the products side of each irreversible reaction (Fig 3.A). Consequently, Eq 19 would no longer interfere with the constraints on the selection of *K*s for the products. This auxiliary species acts as a confiner on its own by following the same assumption applied in Eq 19 but without putting any extra constraints on the products (Eq 20). *K*_aux_ in Eq 20 corresponds to the auxiliary species’ constant.

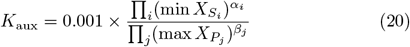 The rest of the process is similar to the reversible reactions in which *K*s are estimated by generating and solving a linear matrix and will be used to approximate the reaction rate constants during the data fitting.
- **Irreversible Michaelis-Menten:** To convert an irreversible Michaelis-Menten reaction into bond graphs, the same configuration was used as in the reversible Michaelis-Menten. Fig 3.B shows an exemplar irreversible Michaelis-Menten reaction. Note that here, reaction 2 is irreversible. Hence, as discussed, the constraint in Eq 19 must be applied. Again, in case of the irreversible Michaelis-Menten reaction being present in a thermodynamically inconsistent loop, an auxiliary species (*C_S_* : *X_aux_*) with *K_aux_* gained from Eq 20 will be added to the right side of the reaction. To calculate the equivalent bond graph parameters we applied the technique introduced in [37]. Briefly, a complex enzymatic reaction in Marcelin–de Donder kinetics can be converted into the irreversible Michaelis-Menten kinetics using some assumptions. These assumptions along with estimating the irreversible Michaelis-Menten parameters from the simulation data provide the grounds to estimate the equivalent bond graph parameters. Eq 21 shows the irreversible Michaelis-Menten kinetic law where v is the reaction rate, *X_S_* is the substrate concentration, *V_m_* is the highest molar flow rate in the reaction, and *K_m_* is the substrate concentration where the reaction rate is half its highest value.

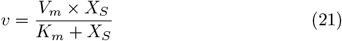 The bond graph form with Marcelin-de Donder parameters in Fig 3.B will give:

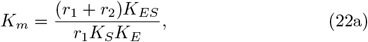

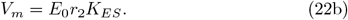

where *E*_0_ is the total concentration of the enzyme and enzyme complex: *E*_0_ = *X_E_* + *X_ES_*. Finding bond graph parameters from the irreversible Michaelis-Menten rate law in Appendix C elucidates the detailed procedure. By having *K_m_* and *V_m_* as constants and applying Eq 19 to implement irreversible reactions, a solution with the least square error can be achieved for the unknown bond graph parameters (*r*_1_, *r*_2_, *K_S_*, *K_P_*, *K_ES_*, *K_E_*, *E*_0_).

To more comprehensively demonstrate the features of our implementation, we created an SBML model of four reactions, including reversible and irreversible mass action kinetics and reversible and irreversible Michaelis-Menten kinetics. The automatically converted model in bond graphs is deposited on Github: [Reversible/Irreversible Mass action and Michaelis-Menten](https://github.com/Niloofar-Sh/SBMLtoBGs/blob/main/reversibleMA_reversibleMM_irreversibleMA_irreversibleMM/reversMAMM_irreversMMMA.ipynb).

In the next section we explain our workflow in dealing with single or multiple SBML models.

### 2.3. The workflow

This section describes the methodology for converting SBML models into bond graph approximations. If we intend to compose the converted models later, we need to estimate the bond graph parameters for the whole composed system. Both cases are discussed in this section.

In brief, the bond graph parameters of a system are approximated from the simulation data during the time course where the entities reach their steady-state behaviours. We collected this data by running the models in SBMLsimulator [38, 39] (www.ra.cs.uni-tuebingen.de/software/SBMLsimulator) and saving the simulations as csv files. The bonds between the bond graph components are defined by the reactant(s)-reaction-product(s) relationships within the SBML models. These information along with the models’ metadata were read using two Python libraries: libSBML [40] and simpleSBML [41]. Because SBML modelling packages and programs do not include the bond graph structure, we have used a bond graph library in Python (BondGraphTools [27]) to construct our bond graph models. To automatically convert an SBML model into bond graphs using our framework, the original model must meet the following criteria:

1. The reactions must be described in terms of one of the following rate laws: Marcelin-de Donder kinetics (using thermodynamic parameters), reversible/irreversible mass action kinetics, or reversible/irreversible Michaelis-Menten kinetics;
2. The model must not contain modifiers. A modifier is a molecule that locally influences the conformation of a component [42]. Modifiers can represent various structures, compounds, and states such as ligands, covalent bondings, and mutations.
3. The model only contains constant parameters;
4. Some SBML models incorporate “events”, which are discontinuous changes in models triggered by certain conditions or at certain times [43]. Since these events are not a part of physical or biological systems, we do not include them in our conversion framework.

If a reaction is formulated with a more complex rate law that does not fit in any of the mentioned categories in 1, as a final attempt, our framework tries to approximate it with reversible/irreversible mass action kinetics. However, this may or may not provide an accurate approximation of the original model. To estimate the number of SBML models compatible with our approach, we considered the first 200 SBML models on BioModels (out of over 2000 available) and 35 of them complied with the mentioned criteria. Therefore, we expect that hundreds of convertible models exist on the Biomodels database (with more than 2000 curated and non-curated models).

Fig 4 illustrates our workflow to create bond graph equivalents of single SBML models of biochemical pathways. The whole workflow can be described in eight major steps as follows:

1. The simulation data are collected by running an SBML model under the model’s initial conditions and saving the results in csv format.
2. We have designed the framework to extract and modify the concentration units to *μ*mol if required. This could be set to any unit, but we selected *μ*mol as it seems to be the most commonly used unit in modelling metabolic pathways on BioModels. The user can skip this step if a single SBML model is the target of the conversion. The user can also insert a scaling factor multiplied by all the concentrations (the default value is 1.0). As discussed later in Section 3.3, the scaling factor is valuable for model integration where the concentrations need to be made consistent between models.
3. In this step, the reaction types (reversibility/irreversibility) and rate laws are inferred from the model’s metadata. Our framework extracts and categorises the reactions’ data by identifying their Systems Biology Ontology (SBO) terms [44].
4. The bond graph parameters (species and reaction rate constants) are calculated in this step. The reaction specific constraints are inferred from the simulation data to calculate the species constants (Section 2.2.1 & Section 2.2.2).
5. This step checks the consistency of semantic annotations within an SBML model. A function in our framework looks for the existence of duplicate annotations of species within the SBML model. If two or more species are annotated identically in one compartment, our function assumes they refer to the same entity and merges them.
6. The bond graph structure of the model (without assigning numerical parameter values) is generated from the stoichiometry of the SBML model. We call these symbolic models.
7. BondGraphTools generates the ODE equations based on the bond graph graphical structure from step 6.
8. In our workflow, the calculated bond graph parameters in step 4 parameterise the generated ODE equations in step 7.

**Fig 4.**
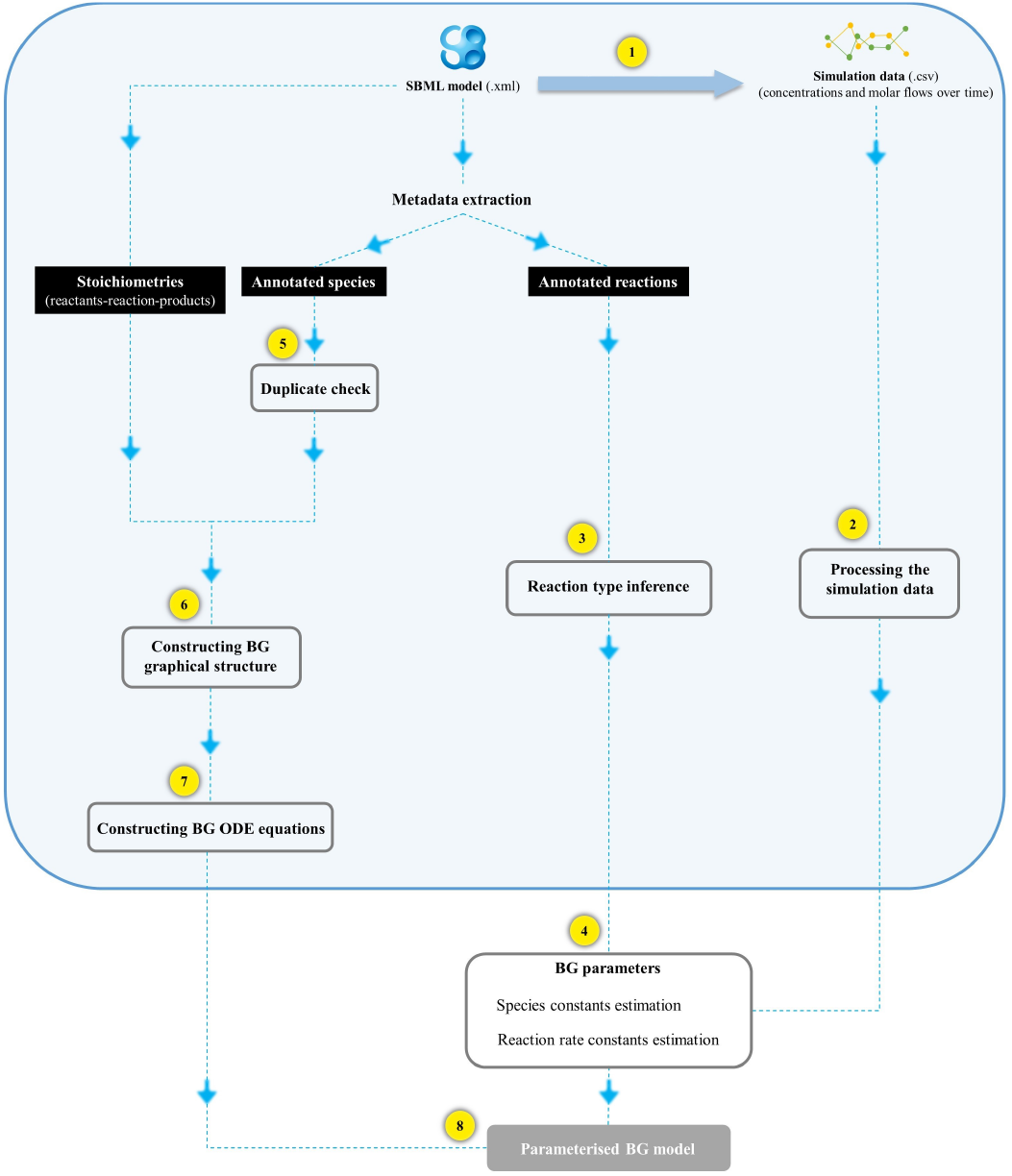
Workflow to generate the equivalent bond graph model of an SBML model. The workflow comprises of eight main steps where the first four steps are involved in estimating the bond graph parameters, and the final three steps generate the bond graph graphical structure and ODE equations. The blue box shows the same procedure performed during model composition and will create a module for each model.

In the case of model composition, the steps in the blue box in Fig 4 remain in place and create a module per SBML model. Each module will have two sets of outputs: 1) An equivalent symbolic bond graph model and 2) the classified constraints and the (unified) simulation data. Fig 5 demonstrates the simultaneous creation of a composed symbolic bond graph model of two modules and the estimation of bond graph parameters. To deal with potential model inconsistencies, when dealing with more than one SBML model, the constraints for all the models must be considered to calculate the bond graph parameters (step 5). Thus, we require that the estimation of bond graph parameters and the creation of bond graph symbolic structure take place out of the modules to include all the information from all the modules. Similar to the final procedure in Fig 4, the species and reaction rate constants are used to parameterise the composed symbolic bond graph model. It is worth mentioning that in cases where the initial condition of the same species is different among the models, our function gives the user the option of choosing any arbitrary amount.

**Fig 5.**
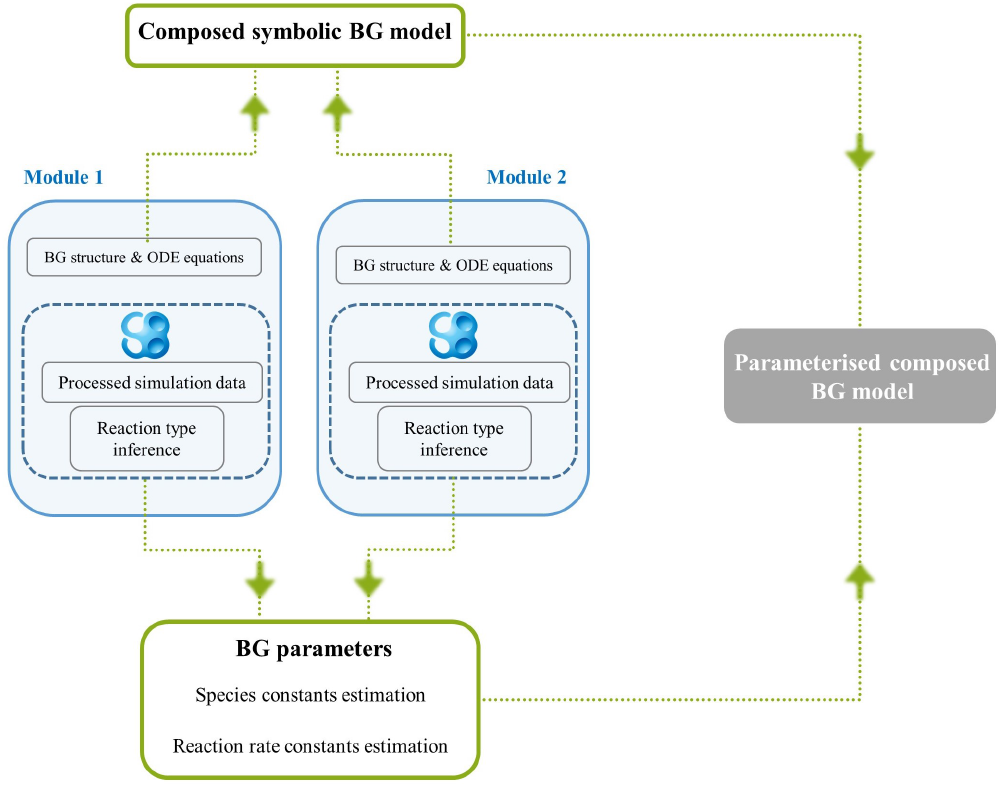
Modular model composition of bond graph equivalents of SBML models. The workflow is drawn for two modules but is extendable to any number of modules. The bond graph parameters and symbolic model are determined by including all the constraints of the selected SBML models.

### 2.4. SBML models

In this section, we introduce the two SBML models (pyruvate distribution and PPP) that we selected for automatic conversion into bond graphs using the framework described in Section 2.3. We selected these two models because, first, they meet the criteria mentioned in Section 2 and second, they have common species, which is required later for composing the models. Here, we review the function and components of each model.

#### 2.4.1. First model: pyruvate distribution

The first selected model is BIOMD0000000017 from the BioModels repository. It describes lumped glycolysis plus branches around pyruvate metabolism in lactic acid bacteria, developed by Hoefnagel et al. [45]. Glycolysis occurs in the cytosol where glucose oxidises to pyruvate [46]. Pyruvate undergoes several processes and produces acetyl CoA through one of them. Fig 6 illustrates the network and the involving biochemical species in the pyruvate distribution model. Purple species represent the ones which are common with the second model (PPP). Table 2 gives the definition for the abbreviations used in Fig 6.

**Table 2.** Abbreviations in the first model: pyruvate distribution.

| Abbreviation | Definition |
| --- | --- |
| *Ac* | Acetate |
| *AcCoA* | Acetyl CoA |
| *AcLac* | Acetolactate |
| *AcO* | Acetaldehyde |
| *AcP* | Acetyl phosphate |
| *ADP* | Adenosine diphosphate |
| *ATP* | Adenosine triphosphate |
| *CoA* | Coenzyme A |
| *EtOH* | Ethanol |
| *NAD* | Nicotinamide adenine dinucleotide |
| *NADH* | Reduced nicotinamide adenine dinucleotide |
| *O<sub>2</sub>* | Oxygen |
| *P<sub>i</sub>* | Phosphate |

**Fig 6.**
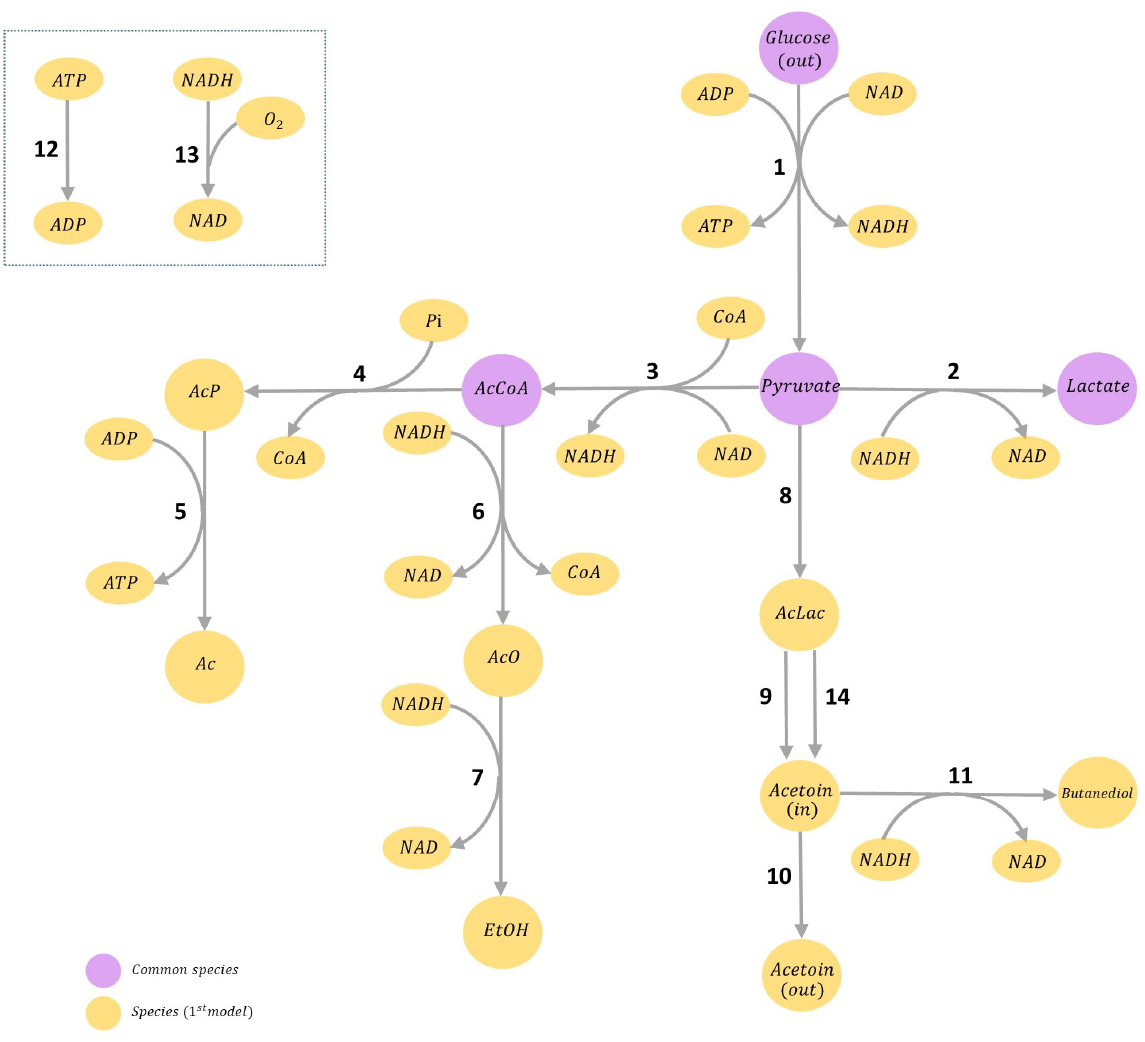
Pyruvate distribution model: Lumped glycolysis and branches around pyruvate metabolism. The species in purple are common between the pyruvate distribution (first) model and PPP (second) model. (out) and (in) refer to extracellular and intracellular concentrations, respectively. Reactions 12 and 13 in the dashed box are common between multiple steps. (Online version in colour.)

The reactions in the pyruvate distribution model are all modelled irreversibly in BIOMD0000000017, but the rate laws are neither mass action nor Michaelis-Menten. Nevertheless, our framework managed to approximate them as bond graphs using the irreversible mass action kinetics (method explained in Section 2.2.2).

#### 2.4.2. Second model: PPP

The second selected model is MODEL1004070000 from BioModels. It describes the pentose phosphate pathway and TCA cycle in the lactating rat mammary gland, developed by Haut et al. [47]. Physiologically, the pentose phosphate pathway is parallel to glycolysis, and they share some reactions. In the PPP model, acetyl CoA enters the TCA cycle [46]. The TCA cycle is a respiratory process that occurs in the cytosol and generates citric acid. The TCA cycle in the current model is represented crudely and hence, all the compounds with six molecules of carbon are collectively referred to as *C6 comps* (including citric acid). Fig 7 demonstrates the network of the PPP model and its species. The species in purple are shared with the pyruvate distribution (first) model. Table 3 gives the definition for the abbreviations used in Fig 7.

**Table 3.**
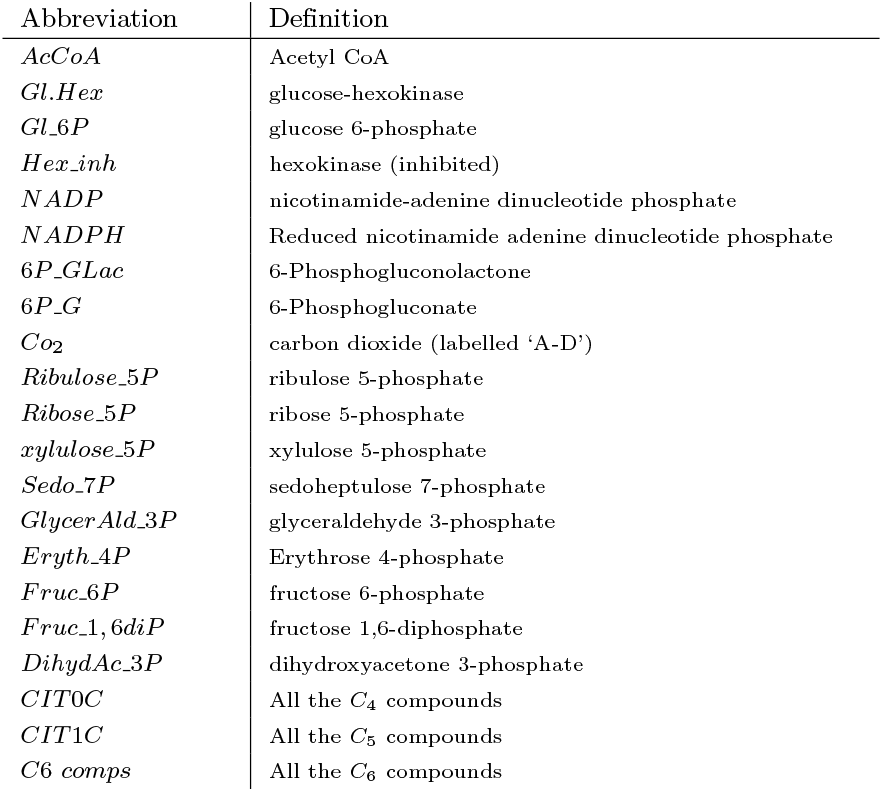
Abbreviations in the second model: PPP.

**Fig 7.**
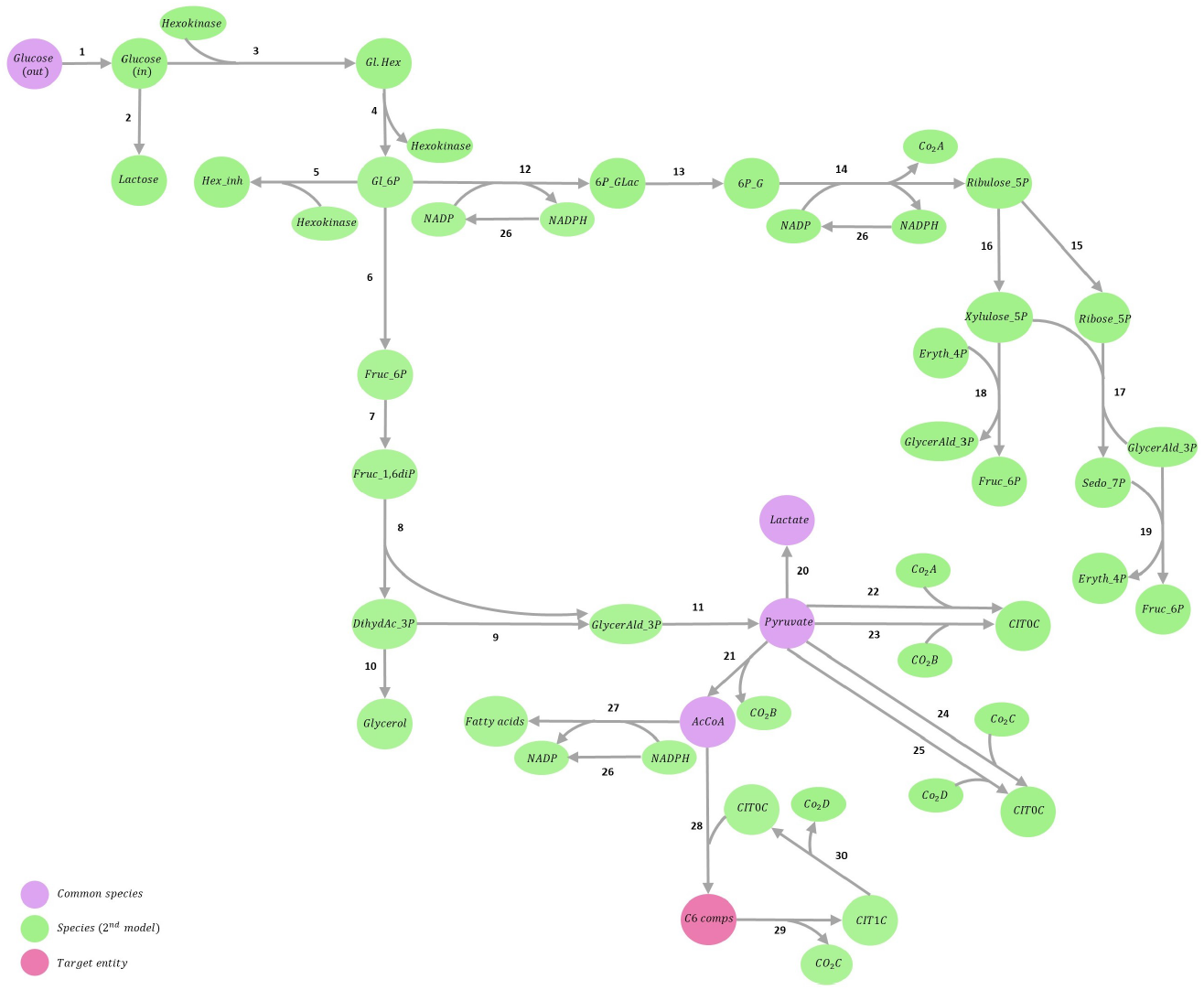
PPP model: pentose phosphate pathway and lumped TCA cycle. The species in purple are shared between the pyruvate distribution (first) model and PPP (second) model. *C6 comps* (in red) corresponds to all the compounds in the pathway with 6 carbon molecules (referred to as the target entity), including citric acid. (out) & (in) refer to extracellular and intracellular concentrations, respectively. (Online version in colour)

Reactions in the PPP model are a combination of 19 reversible and 11 irreversible reactions, and all the reactions are defined in mass action kinetics. Thus, using the methods described in Sections 2.2.1 & 2.2.2, the equivalent bond graph model was created and parameterised.

## 3. Results

We used our developed framework to automatically convert two exemplar SBML models into bond graphs. We verified the simulations by comparing them with the original models to validate our equivalent bond graph models. To compare the simulation results, the normalised root mean square error (NRMSE) was computed as in Eq 23, where 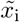 corresponds to the simulation points of the bond graph approximation, and *x_i_* corresponds to the original SBML model. The normalisation was performed relative to the maximum and minimum data difference from the SBML model in each simulation.

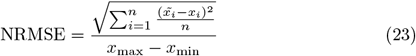

### 3.1. Pyruvate distribution model approximation in bond graphs

The responses of six exemplar species in the pathway are demonstrated in Fig 8. Four of these species are common in the two selected SBML models (Glucose, Lactate, AcCoA, and pyruvate). Glucose, Lactate, Ac, EtOH, Acetoin (out), Butanediol, O2, and PO4 are chemostats in this model. The NRMSE is computed for each comparison in percentage.

**Fig 8.**
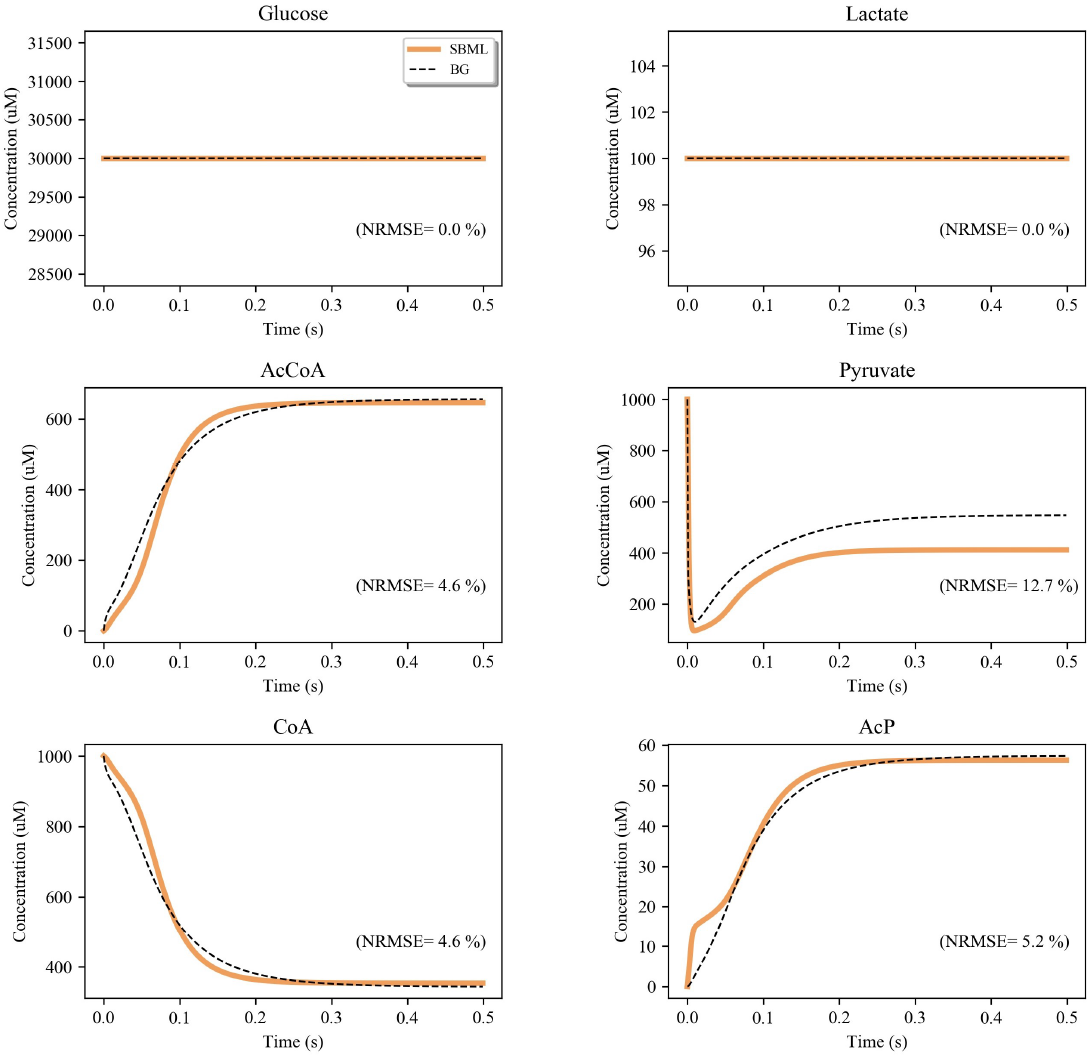
Comparison between the pyruvate distribution SBML model and its bond graph (BG) approximation. The simulations are given for six exemplar species in the pathway. NRMSE is calculated for each comparison in percentage.

As noted earlier, an exact conversion from SBML to a bond graph was not possible due to the approximation of the irreversible reactions with reversible reactions. Nonetheless, the generated bond graph model exhibits similar behaviour to the original SBML model. Note that none of the rate laws in this model were described using the reversible/irreversible mass action or Michaelis-Menten. In such cases our function tries to approximate the reactions in bond graphs by assuming that they follow the mass action kinetics. This approximation resulted in a feasible bond graph approximation of the SBML model despite the most noticeable difference of 12.7% NRMSE in pyruvate reaching its final state. However, we note that these differences may increase when the models are simulated under different conditions.

### 3.2. PPP model approximation in bond graphs

Fig 9 demonstrates the comparison between the behaviour of the PPP SBML model and its equivalent bond graph approximation.

**Fig 9.**
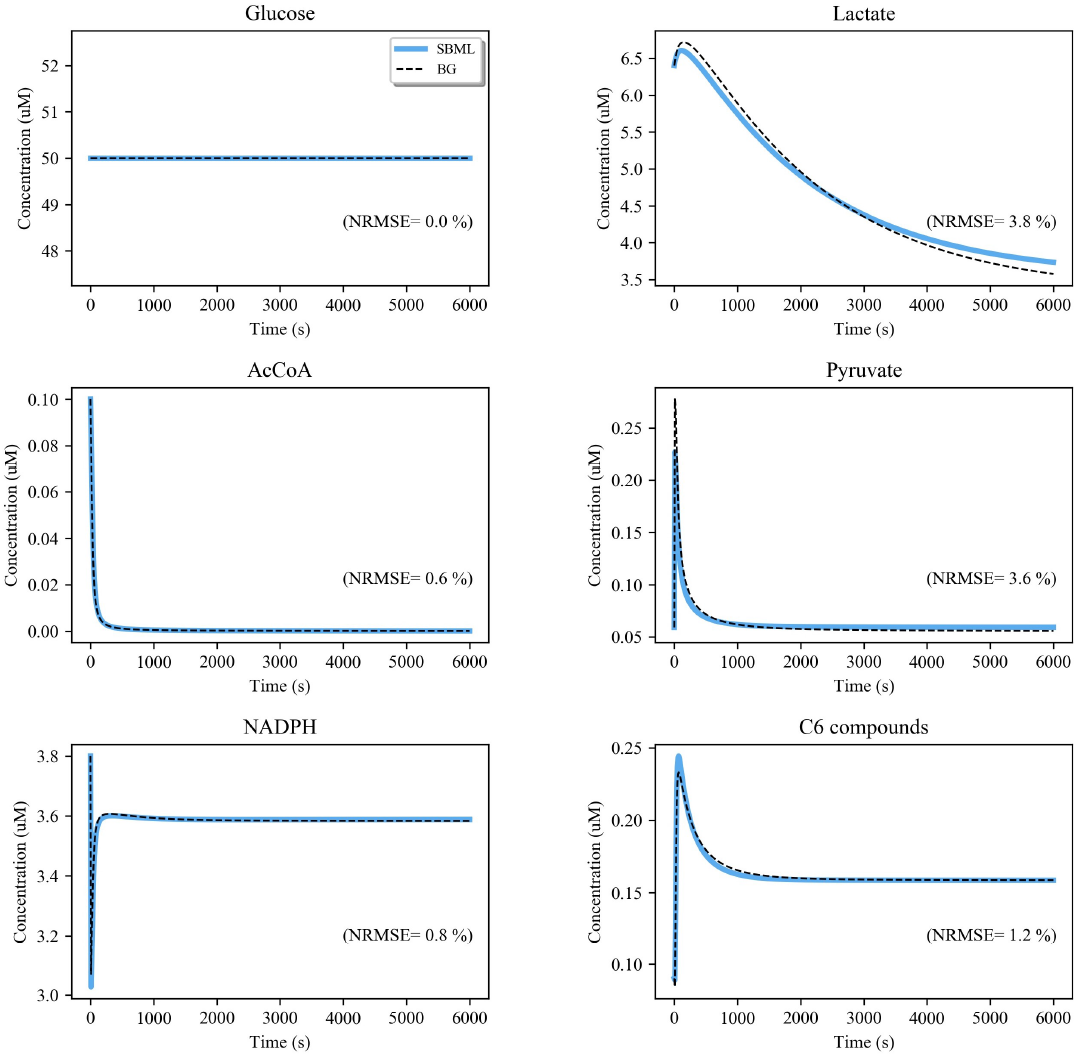
Comparison between the PPP SBML model and its bond graph (BG) approximation. The simulations are given for six exemplar species in the pathway. NRMSE is calculated for each comparison in percentage.

The equivalent bond graph model replicated the transient performance of the original SBML model. As in the first model, an identical performance was not achieved because some kinetic parameters in the original model were not thermodynamically consistent. The approximation of irreversible reactions with reversible ones also led to some minor inconsistencies. Here, the bond graph conversion of the PPP model showed an error of 3.8% and 3.6% NRMSE in replicating the behaviour of lactate and pyruvate, respectively.

### 3.3. Model composition

In this section we demonstrate the application of our automated bond graph model composition methodology to our generated models. The composed model will be thermodynamically consistent and easily reused due to the intrinsic features of bond graphs (Section 1). The automatic composition and merging has become possible through finding identical semantic annotations among models. Liebermeister et al. had previously developed a tool (SemanticSBML) to automatically merge SBML models by matching their annotations [48]. The same idea has been used in our work but instead of matching the original SBML models, we applied this method to the devised bond graph models. Although we merge bond graph modules, the result of the model composition is a single (flattened) bond graph model. We have performed the composition using the approach introduced in [24] with some improvements and added features:

1. **SBML support:** Since the framework was initially developed for CellML models, we slightly modified it to load and extract information from SBML models;
2. **Processed data:** To help couple models together, we added the *units unification* and *scaling* modules to the framework to help with model composition as discussed in Section 2.3, step 2 of the workflow. After unit unification, the user can select a scaling index for any of the models. The scaling index will be multiplied to the concentrations of the selected model. This normalises the amounts where there is a significant difference between the range of amounts in the selected models;
3. **Species constants inconsistencies:** There are cases where the same species within multiple bond graph models have been granted different values for their species constants (depending on the simulations and parameters extracted from each reference model). This causes inconsistencies during model composition since only one value can be assigned to a merged species. The method introduced in [24] dealt with this by asking the user to choose between the values or enter a new value. Here, we improved the model composition framework by inclusively considering all the constraints from the reference SBML models (as discussed in Section 2.3). In this way, only one value is assigned to each species constant and the need for the user decision is lifted;
4. **Removable components:** Users can select desired component(s) to be removed from models. This feature is useful in the case of unidentified duplicates, which can arise due to improper annotation or appearances of the same process in different models.

Here, the pyruvate distribution model corresponds to lactic acid bacteria and the PPP model corresponds to lactating rat mammary glands. As a result, the range of concentrations in the pyruvate distribution model is approximately 1000 times higher than in the PPP model. Thus, we decided to input a scaling index of 0.001 for the former model. Considering the different origins of the constitutive models, having the concentrations in the same range induces more realistic behaviours in the composed model. While merging these two models may not precisely represent an existing biological system, it will result in a physically plausible model that provides the groundwork for the development of more realistic models in the future.

The *Glucose* (*out*) → *Pyruvate* reaction in the pyruvate distribution model lumps together several steps from *Glucose* (*out*) to *Pyruvate* in the PPP model (Fig 10). Therefore, we decided to remove the abstracted step (in the pyruvate distribution model) from the final composed model. Our framework deals with common reactions (here, *Pyruvate → AcCoA* and *Pyruvate → Lactate*) automatically. It uses the information on the reactant(s) and product(s) of each reaction and if they are identical, our framework considers the reactions being duplicates. Thereafter, it keeps one of the reactions and removes the rest.

**Fig 10.**
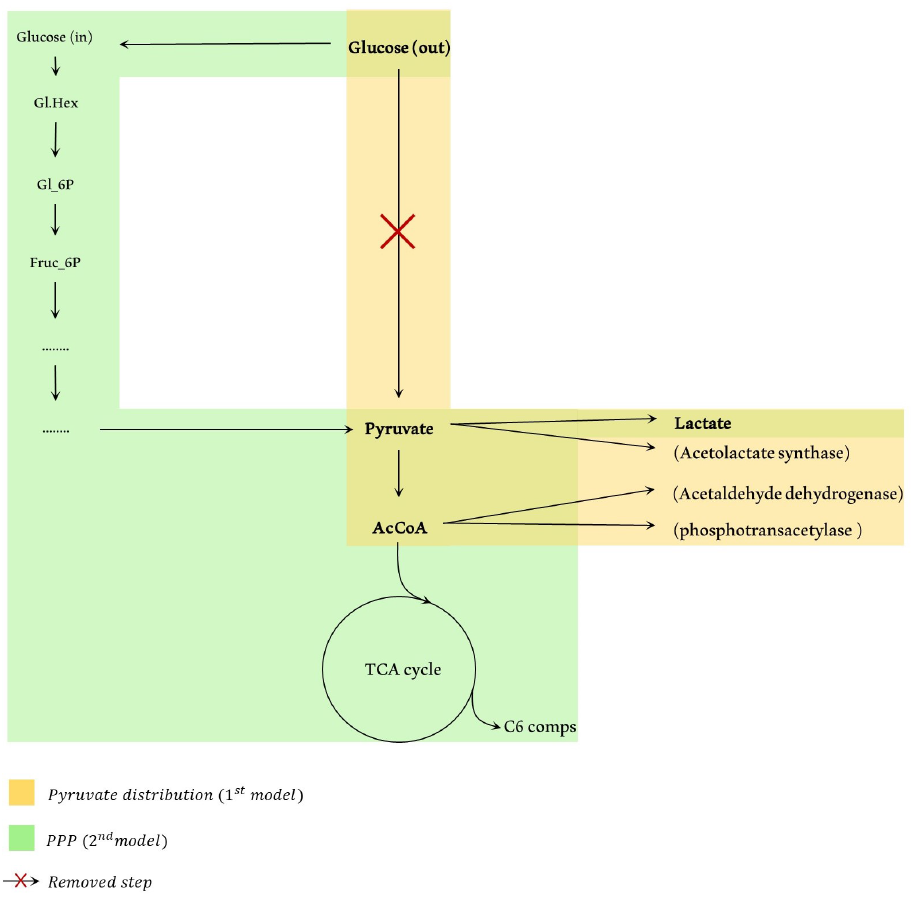
Pyruvate distribution and PPP models with the species in common. Four common species are present in both models: Glucose (out), Pyruvate, AcCoA, and Lactate and the two common reactions are: *Pyruvate* → *AcCoA* and *Pyruvate* → *Lactate*. Subsidiary processes are shown in parentheses for clarity. The manual removal of the *Glucose* (*out*) → *Pyruvate* abstract step from the pyruvate distribution model is specified by a red cross. (Online version in colour)

The PPP and glycolysis are fueled by glucose [49, 50]. To verify the behaviour of the final composed model, we simulated our bond graph “pyruvate distribution+PPP” model for different extracellular concentrations of glucose and monitored how the concentrations of pyruvate and C6 compounds were correlated to this change. Pyruvate is present in both models and C6 compounds are a group of target products (including citric acid) in the PPP model. Fig 11 illustrates the behaviour of pyruvate and C6 compounds under the condition of changing the extracellular concentration of glucose in 6 steps from 25 *μ*M to 500 *μ*M. Fig 11 shows that the higher the concentration of extracellular glucose, the higher the concentration of pyruvate and C6 compounds, implying the role of glucose in initiating and functionality of the composed pathway. The glucose-pyruvate relationship was also studied by Zhu et al. in [51], where altering the glucose levels directly affected the pyruvate concentration. Papagianni et al. [52] also showed that the formation rate of citric acid increases with increasing glucose levels.

**Fig 11.**
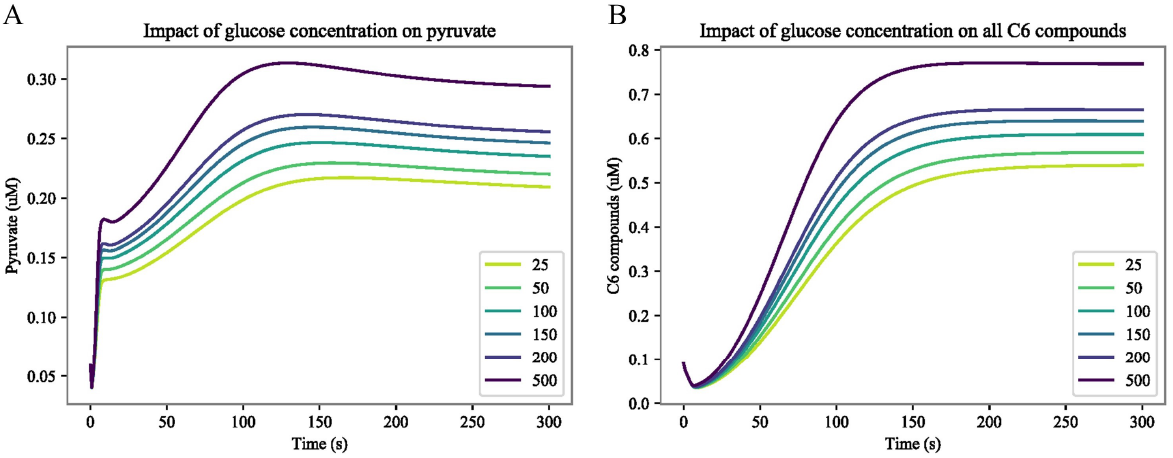
Effect of different levels of the extracellular concentration of glucose on pyruvate and C6 compounds. Legends show the extracellular glucose concentration in 6 steps (25-500 *μ*M). (A) Pyruvate behaviour. The initial concentration of the other common species were as: pyruvate=0.059 *μ*M, AcCoA=0.1 *μ*M, and lactate=100 *μ*M (fixed); (B) C6 compounds behaviour. (Online version in colour.)

## 4. Discussion and conclusion

In this paper, we introduced a new methodology for converting SBML models to bond graphs. This was achievable because the reference models contain annotations for biochemical species and rate laws. The bond graph parameters were calculated from the simulations to approximate the behaviour of the original models. The resulting bond graph models could be automatically composited. We applied this method to convert and merge two SBML models: pyruvate distribution and the pentose phosphate pathway (PPP). Bond graph equivalents of other exemplar SBML models – generated using our function – can be found on GitHub: [other SBML to BG conversions](https://github.com/Niloofar-Sh/SBMLtoBGs/tree/main/Verified_other_conversions).

Our tool is fundamentally different from the SHMC package and semanticSBML. The SHMC package and semanticSBML combine models while retaining their original modelling scheme, which in most cases represents physically impossible models. Also, changes in equations in model compositions must be managed by the user, which is an error-prone and time-consuming task. Our tool avoids these issues by converting the models into bond graphs and merging them on an energy-conserving platform. This automates the SBML→BG conversion which by incorporating annotations, also automates model composition. SBML models that meet the criteria must be fully annotated with added SBO terms for rate law recognition.

It is worth mentioning that the reversible Michaelis-Menten kinetics can be implemented in bond graphs using a single *Re* component with the constitutive equation of the form Eq 16. However, since the constitutive equation of the existing *Re* component in BondGraphTools is described by the Marcelin–de Donder equation, we decided to use the configuration introduced in [20] as a formulation for the enzyme-catalysed reactions.

In bond graph model composition, there are cases where a species is characterised differently in the selected models. The decision to choose either of the types depends on the application. For example, lactate had a fixed concentration in the pyruvate distribution model. So it was modelled as a chemostat (*C_S_*) in bond graphs, while in the PPP model, lactate had variable concentration and was modelled as a normal biochemical entity (*C_e_*). We selected a fixed concentration for lactate while merging the two models, but the users will have both options. In general, three situations might occur during model composition where the user’s decision is required: 1) different initial concentrations of the same species, 2) different definitions of the same species, and 3) adding auxiliary species to irreversible reactions. A modeller’s domain specific knowledge is required to resolve the first two issues. However, the final case could potentially be automated using genome-scale metabolic models (GSMMs). GSMMs are comprehensive stoichiometric networks of metabolic reactions and biochemical data, which help to comprehend the metabolism and physiology of the organisms [53, 54]. These can be employed as scaffolds to automatically detect and locate the missing energy sources and avoiding the need for auxiliary species.

In this paper, we merged models from two different organisms to demonstrate our tool’s capability in generating physically plausible composed models agnostic to any specific underlying biology. Here, we aimed to find SBML models on BioModels to first comply with the four criteria (Section 2) and second, be in the same organisms and third, have common entities to be merged. There were limited choices to fulfil all these benchmarks on a group of about 2000 (curated and non-curated) models on BioModels. Although biologically impossible, we selected the pyruvate distribution of lactic acid bacteria to be merged with the PPP in lactating rat mammary glands to illustrate the possibility of such combinations (useful in examining hypotheses in hybridisation) in an energy-based framework.

In biology, the PPP, glycolysis and the TCA cycle have some more points of connection, namely ADP, ATP, NAD(P), and NAD(P)H. Since NAD and NADH only exist in the first model (pyruvate distribution) and are replaced by NADP and NADPH in the second model (PPP), merging them was not feasible. Moreover, ADP and ATP are not included in the PPP model, which led to some missing connections among the models. While the full reaction network of the pathways in this paper are well known, it is common that a scientist will model a biological system without full knowledge of the chemical reactions. To allow for bond graphs to be generated even in the absence of this knowledge, we decided use general auxiliary species rather than to ask the modeller to include the correct side species themselves. However, as mentioned earlier, GSMMs could be used to help automate this process. This will significantly assist other modellers in reusing pre-existing models in a more physically plausible manner and broader applications.

In this paper, we stored the simulation data for each model as a *csv* file before commencing the conversion process. This can be facilitated in future through the automatic generation and storage of simulations by running the SBML models directly in Python. Also, since we need the species in each model to reach their final concentrations, the running time must be set accordingly. In this paper, we set the running times manually, but in future, automatic recognition of steady-state concentrations can significantly reduce the manual preparation of prerequisites. We also expect that fitting parameters to a wider range of simulation data would allow the generated bond graph models to better match the biology.

Our tool introduces the foundation for making the SBML models thermodynamically consistent and more easily reusable. The present paper is a starting point to facilitate the selection, conversion, and composition of SBML models in an energy-based environment. In future, further investigations can be done to enhance our tool to support the conversion of other standard rate laws found in SBML models such as Hill equation [55], convenience [56], and lin-log kinetics [57] into bond graphs. Similarly, further investigations can be performed to explore the composition of bond graph models from non-SBML sources such as rule-based models [58, 59]. Beyond the SBML models, we also see this is a first step toward making the large number of models that already exist available for maximal reuseability building on the thermodynamic framework that Edmund J Crampin established.

## Appendix A. Matrix equation example

Consider the following two reactions with the kinetic constants 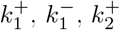, and 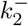.

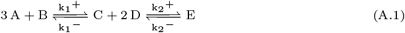

where:

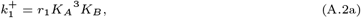

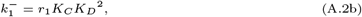

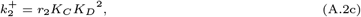

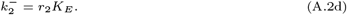

Taking logarithms from each equation gives a matrix of linear relationships:

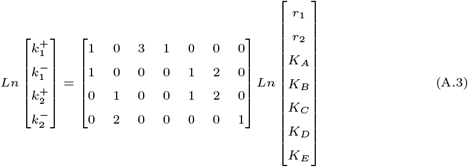

In cases where *k*^+^ and *k*^−^ are not properly annotated and hence not specifically detectable, each reaction will provide us with only one constraint (instead of two for *k*^+^ and *k*^−^). As discussed in Section 2.2.1, we will have access to the ratios of *K*s between the reactants and products of each reaction. Here, *σ*_1_ and *σ*_2_ are the constant ratios for each reaction (gained from the simulation data):

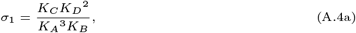

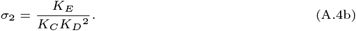

The linear equation matrix for Eqs A.4a and A.4b will be as follows:

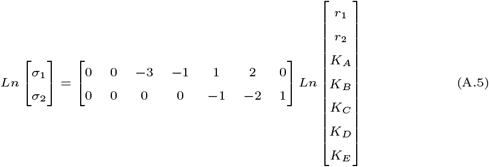

In such cases, the reaction rates (*r*_1_ and *r*_2_) will be determined by curve fitting methods once the equation matrix is solved.

## Appendix B. Reversible Michaelis-Menten

The ratios between the four extracted constants from the reversible Michaelis-Menten formula give the relationships between *K_P_*, *K_S_*, *r*_1_, and *r*_2_:

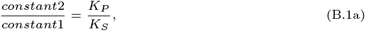

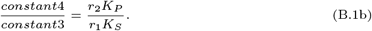

When the matrix of constraints is solved and the values for *K_P_*, *K_S_*, *r*_1_, and *r*_2_ are estimated, the remaining parameters (*K_E_* and *K_ES_*) can be calculated.

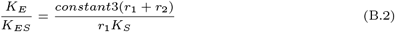

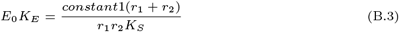

By selecting any arbitrary value for *E*_0_ and substituting Eq B.3 in Eq B.2, we can calculate *K_E_* and *K_ES_* from the known values. Here, we selected *E*_0_ = 1.

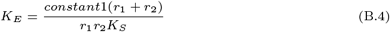

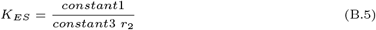

## Appendix C. Irreversible Michaelis-Menten

Here, we demonstrate the Marcelin-de Donder equations for Fig 3.B can be reduced to the irreversible Michaelis-Menten configuration with simplifying kinetic assumptions [60]. We initially illustrate the calculations without the participation of *C_S_* : *X_aux_* and later how a slight change will include the auxiliary species in the equations. The fluxes of the substrate and product (*v_S_* and *v_P_*) reads:

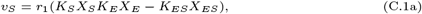

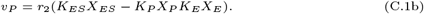

The kinetic equations for the fluxes of *E* and *ES* are given in Eqs C.2a and C.2b which conclude a constant amount of the enzymes in Eq C.2e.

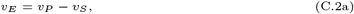

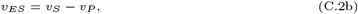

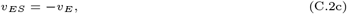

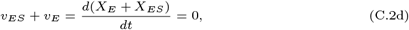

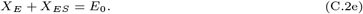

Assuming that the concentration of *E* and *ES* will be constant at steady state, by substituting *v_E_* =0 and *v_ES_* = 0 in Eqs C.2a and C.2b, we have:

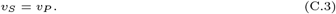

According to the definition of an irreversible Michaelis-Menten, *Re* : *r*_2_ only proceeds in forward direction. This implies that in its reversible mass action configuration, the forward flux must be much greater than the reverse flux. We implement this assumption by selecting *K_P_* ≃ 0 and Substituting Eqs C.1a and C.1b in Eq C.3:

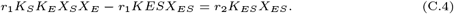

Replacing Eq C.2e in Eq C.4 will read:

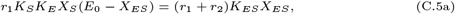

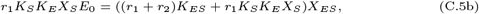

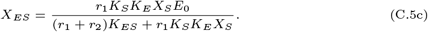

Since *v_P_* = *r*_2_*K_ES_X_ES_* (due to the pseudo-irreversibility of *Re* : *r*_2_), substituting Eq C.5c will give:

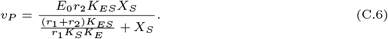

Eq C.6 is the irreversible Michaelis-Menten rate law in Eq 21 where *K_m_* = (*r*_1_ + *r*_2_)*K_ES_*/(*r*_1_*K_S_K_E_*) and *V_m_* = *E*_0_*r*_2_*K_ES_*.

To estimate the parameters of the irreversible Michaelis-Menten kinetics in bond graphs we need to make an initial guess to limit the solutions. Here, we initialised *E*_0_ = 1 which led to the following constraint:

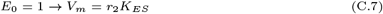

By calculating *r*_2_ and *K_ES_* from the linearised matrix of constraints and substituting the values in the *K_m_* formula, we obtain a relationship between *r*_1_ and *K_E_* as in Eq C.8. We can select any arbitrary value for either *r*_1_ or *K_E_* and calculate the other parameter from Eq C.8. Here, we selected *r*_1_ = 1 and calculated *K_E_*, accordingly (Eq C.9).

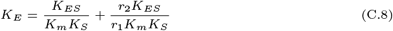

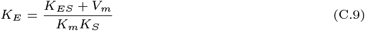

## Notes

https://github.com/Niloofar-Sh/SBMLtoBGs

### Competing Interest Statement

The authors have declared no competing interest.

